# Spatiotemporal clustering of highly pathogenic avian influenza (HPAI) H5N1 at the wild waterfowl-poultry interface: Vector-specific spillover risks in the U.S., 2022–2025

**DOI:** 10.64898/2026.03.06.710020

**Authors:** Csaba Varga

**Affiliations:** University of Illinois Urbana-Champaign, College of Veterinary Medicine, Urbana, Illinois, USA

**Keywords:** Avian Influenza, Poultry, Waterfowl, Infections, space-time cluster, outbreaks

## Abstract

**Background:** The emergence of the highly pathogenic avian influenza (HPAI) H5N1 clade 2.3.4.4b in North America, beginning in February 2022, has highlighted the dynamic, unpredictable, and regionally variable risk of infections. Studies are needed to assess the spatiotemporal clustering of HPAI H5 at the interface between wild waterfowl and commercial poultry to better understand and mitigate this risk.

**Methods:** Publicly available data on HPAI H5 detections in wild birds and commercial poultry from January 2022 to January 2026 were analyzed at the county level. Retrospective space-time permutation models were used to identify and scan for clusters with higher than expected detection rates.

**Results:** A total of 17,091 HPAI H5 detections were reported in wild birds across 1,467 county-level locations. Four species, Mallard (*Anas platyrhynchos*) (2,848 detections, 16.66%), Canada goose (*Branta canadensis*) (1,496, 8.75%), Green-winged teal (*Anas carolinensis*) (1,364, 7.98%), and Snow goose (*Anser caerulescens*) (1,084, 6.34%), accounted for 39.73% of detections. In commercial poultry, 532 outbreaks in turkey operations, 148 outbreaks in table-egg layer operations, 99 outbreaks in broiler chicken operations, and 89 outbreaks in commercial duck operations were reported, respectively. Several spillover events followed an east-to-west expansion. In early 2022, mallard detections preceded outbreaks in Northeast egg-layer and duck farms, while snow goose detections in the Upper Midwest coincided with turkey farm outbreaks. In the Pacific and Mountain West during summer 2022, detections in Canada geese overlapped with turkey farm outbreaks. A resurgence occurred in the Midwest (2025), with snow and Canada goose detections overlapping severe outbreaks in turkey and layer flocks. Additionally, in the Upper Midwest, Canada goose and mallard detections overlapped with outbreaks in commercial duck farms during fall-winter 2025.

**Conclusions:** The study findings demonstrate distinct vector-based transmission dynamics of HPAI H5 at the wild waterfowl-poultry interface. Farm biosecurity strategies must adapt to these recurrent, vector-specific risks.

## Introduction

Avian influenza is an influenza Type A virus, which is classified into different strains based on two viral surface proteins: hemagglutinin (H) and neuraminidase (N). In poultry, highly pathogenic avian influenza (HPAI) viruses are almost always of the H5 or H7 hemagglutinin types, and they can cause severe morbidity and mortality, as well as severe economic concerns [1]. Avian influenza can undergo genetic drift and shift, forming new viral strains, and allowing it to adapt to new host species [2]. Wild waterfowl are considered natural hosts for the virus and can transmit it via migratory flyways, posing a risk to commercial poultry operations [3,4].

The emergence of HPAI strains in North America, specifically the H5N1 clade 2.3.4.4b., beginning in February of 2022, has reinforced how dynamic, unpredictable, and regionally variable the HPAI risk is [5]. Since early 2022, HPAI detections have been confirmed in commercial and backyard poultry flocks, wild waterfowl, wild mammals, and dairy cattle across multiple U.S. states [6,7]. Since the start of the HPAI H5 outbreak in commercial and backyard poultry flocks on February 8^th^, 2022, depopulation of over 195 million birds in the U.S. has been reported to date [8].

Unlike previous HPAI viral strain introductions into the U.S. via migratory flyways, which were restricted to seasonal waves, the current lineage has established endemicity in wild waterfowl populations across the U.S., posing a continuous risk to commercial poultry farms (3). The current HPAI H5 strains show heterogeneity in vector competence and migratory behavior, are stable in the environment, have expanded their host tropism, including spillover into mammals, and can cause die-offs in waterfowl [9].

Current HPAI surveillance systems in the U.S. operate in silos, with data fragmented across monitoring data sets from wild birds [10] and commercial poultry operations [8]. These data silos prevent the generation of cross-sectoral models and real-time risk forecasts that could guide proactive decision-making and early interventions. Moreover, the spatiotemporal transmission dynamics among specific waterfowl species and different commercial poultry sectors are underdefined. Current surveillance often aggregates “wild birds” into a single risk variable, does not account for the distinct ecological roles that each wild waterfowl species plays, nor does it distinguish between the HPAI H5 transmission risk of long-distance migrants versus synanthropic bridge species [11,12].

To address this gap, we integrated HPAI surveillance data from January 2022 through December 2025, obtained from wild birds and commercial poultry operations, to characterize the interface between four key waterfowl vectors, Mallard (*Anas platyrhynchos*), Canada goose (*Branta canadensis*), Snow goose (*Anser caerulescens*), and Green-winged teal (*Anas carolinensis*), and four commercial poultry production types; Turkey, Table egg layer, Broiler, and Duck. We used disease mapping and space-time permutation models to identify areas and time periods with the highest risk of detecting spillover events among wild waterfowl and commercial poultry farms. The final goal of this study is to provide information to animal health stakeholders to aid them in designing early warning systems and location, and species-specific biosecurity plans to mitigate the burden and risk of HPAI H5 in poultry operations across the United States of America (U.S.).

## Materials and Methods

### Data Source

Wild bird HPAI surveillance is overseen by the United States Department of Agriculture (USDA), Animal and Plant Health Inspection Service (APHIS), which collects data on laboratory-confirmed HPAI H5 detections submitted by federal, state, tribal, and partner agencies across the United States. The samples are collected nationwide to represent the national status of HPAI H5 and are obtained through the investigation of morbidity and mortality events, surveillance in live wild birds, hunter-harvested birds, use of sentinel species, and environmental sampling [10,13]. In commercial poultry, HPAI is a nationally reportable disease, and each confirmed and suspected case must be reported to APHIS and State animal health officials [14].

Publicly available data on HPAI H5 detections in wild birds [15] and commercial poultry [8], between January 1^st,^ 2022, and January 9^th,^ 2026, for wild birds and February 08 2022, and January 16^th^, 2026, for commercial poultry, were obtained from the APHIS HPAI detection dashboard. Separate downloadable datasets for wild birds and commercial poultry were retrieved in CSV format and used for subsequent analyses. The data included information on detection date, state, county, species (wild birds), and poultry production type (commercial broiler, layer, turkey, ducks).

### Statistical Analysis

The R software (Version 4.5.2) [16] and the RStudio (Version 2026.01) platform were used for data management and descriptive statistics. Disease maps and spatial statistical analyses were conducted in ArcGIS Pro 10.7.1 (Environmental Systems Research Institute, Inc., Redlands, CA, USA).

The HPAI H5 detections were analyzed at the county level. For each county centroid, latitude and longitude were calculated and represented as point features for spatial analysis. The analysis was restricted to counties within the contiguous U.S. (48 states and the District of Columbia, excluding Alaska and Hawaii) to comply with distance-based analyses. The data was projected to the NAD 1983 (2011) USA Contiguous Albers Equal Area Conic coordinate system to ensure accurate distance calculations.

For each wild waterfowl species and poultry operations, choropleth point maps were constructed using divergent colors and natural breaks (Jenks) classification with 5 categories. Inverse distance weighted interpolation was used to complement choropleth mapping of wild waterfowl HPAI H5 detections and to illustrate the spatial intensity of detections and estimate patterns in areas without reported data.

Retrospective space-time analysis was conducted, scanning for clusters with high rates using the space-time permutation model in the SaTScan software (Version 9.6) [17]. This model does not require population-at-risk data, and it was appropriate for our study because information on the total number of tests performed was unavailable. A cylindrical scanning window was applied, where the circular base represented the geographic area and the height represented the time interval. The maximum spatial cluster size was set to 50% of the population at risk, the minimum temporal cluster size was defined as 1 month, and the maximum temporal cluster size was set to 50% of the study period. Statistical significance (p ≤ 0.05) was assessed using Monte Carlo hypothesis testing with 999 replications. The observed-to-expected (O/E) ratio was calculated for each space-time cluster to quantify excess risk, with values greater than 1 indicating a higher-than-expected number of detections within the cluster window under the null hypothesis of random space-time distribution. All statistically significant space-time clusters were visualized in maps.

## Results

### Distribution of EA H5 HPAI detections in wild birds

Between January 1, 2022, and January 09, 2026, 17,091 laboratory-confirmed detections of EA H5 HPAI were reported in wild birds across 1,467 county-level locations in 48 U.S. contiguous states and the District of Columbia. Four species, Mallard (*Anas platyrhynchos*) (2,848 detections, 16.66%), Canada goose (*Branta canadensis*) (1,496, 8.75%), Green-winged teal (*Anas carolinensis*) (1,364, 7.98%), and Snow goose (*Anser caerulescens*) (1,084, 6.34%), accounted for 39.73% of detections. These four wild bird species were chosen for further analysis due to their detection frequency, broad geographic distribution, and significant roles in avian influenza ecology and transmission to commercial poultry (**Figure 1**).

**Figure 1.**
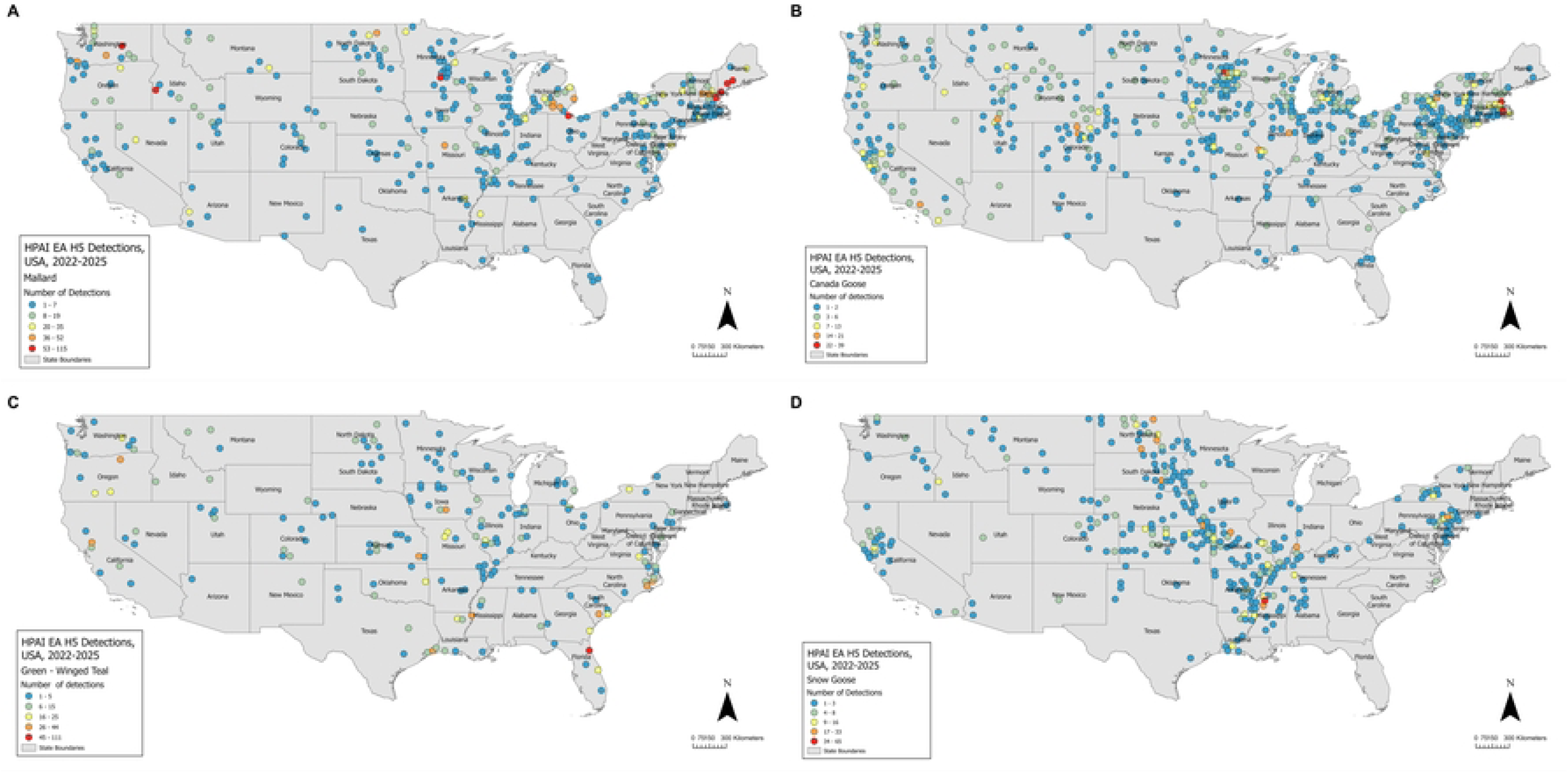
Distribution of highly pathogenic avian influenza (HPAI) H5 detections across the U.S. by wild waterfowl species, 2022-2025. The map highlights detections in four wild waterfowl species: (A) Mallard, (B) Canada goose, (C) Green-winged teal, and (D) Snow Goose. Divergent colors are used, with red indicating high and blue indicating low detection areas.

Mallard detections were distributed across 305 locations, with high concentrations in the Pacific Northwest, Upper Midwest, Great Plains, and Northeast. Canada geese were detected across 500 locations, with high numbers in the Northeast, Upper Midwest, Pacific Coast, and Interior West. Green-winged teal detections were concentrated in the Southeast, with moderate detections across the Pacific Northwest, Pacific Coast, South, and Great Plains. Snow geese were found across 314 locations, with high detections in the South and moderate detections in the Midwest, Great Plains, and Northeast.

### Interpolated HPAI H5 detection intensity across waterfowl species

Spatial interpolation identified distinct high-intensity HPAI detection zones (“hotspots”) that varied by vector species (**Figure 2**). Mallard detections were concentrated in the Northeast (New York, New England) and additional elevated areas in the Upper Midwest (Minnesota, Wisconsin, Michigan) and the Pacific Northwest (western Washington, Oregon). Low intensities were widespread across the Intermountain West, Southwest, and the central and southern Great Plains. High detection intensities for Canada goose. were centered in the Upper Midwest (Minnesota, Wisconsin, Michigan) with additional elevated areas in the Northeast (New York, New England) and the Pacific Coast (California). Low intensities dominated the Intermountain West, Southwest, and much of the central and southern Great Plains. The highest intensities for Green-winged teal detections were concentrated in the Southeast, almost entirely in Florida. Moderate-to-elevated values extended across the South (Louisiana, Mississippi, Arkansas, Missouri) and the Pacific Northwest (Oregon, Washington). Low intensities were widespread across the northern and central United States, including the Upper Midwest, Northern Great Plains, and Intermountain West. High intensities for Snow goose were centered in the South, particularly Mississippi, with elevated zones extending through the Central Midwest and Northern Great Plains (Missouri, North Dakota, South Dakota) and into the Northeast (Pennsylvania). Low intensities were broadly distributed across the Intermountain West, Southwest, and large portions of the Great Plains.

**Figure 2.**
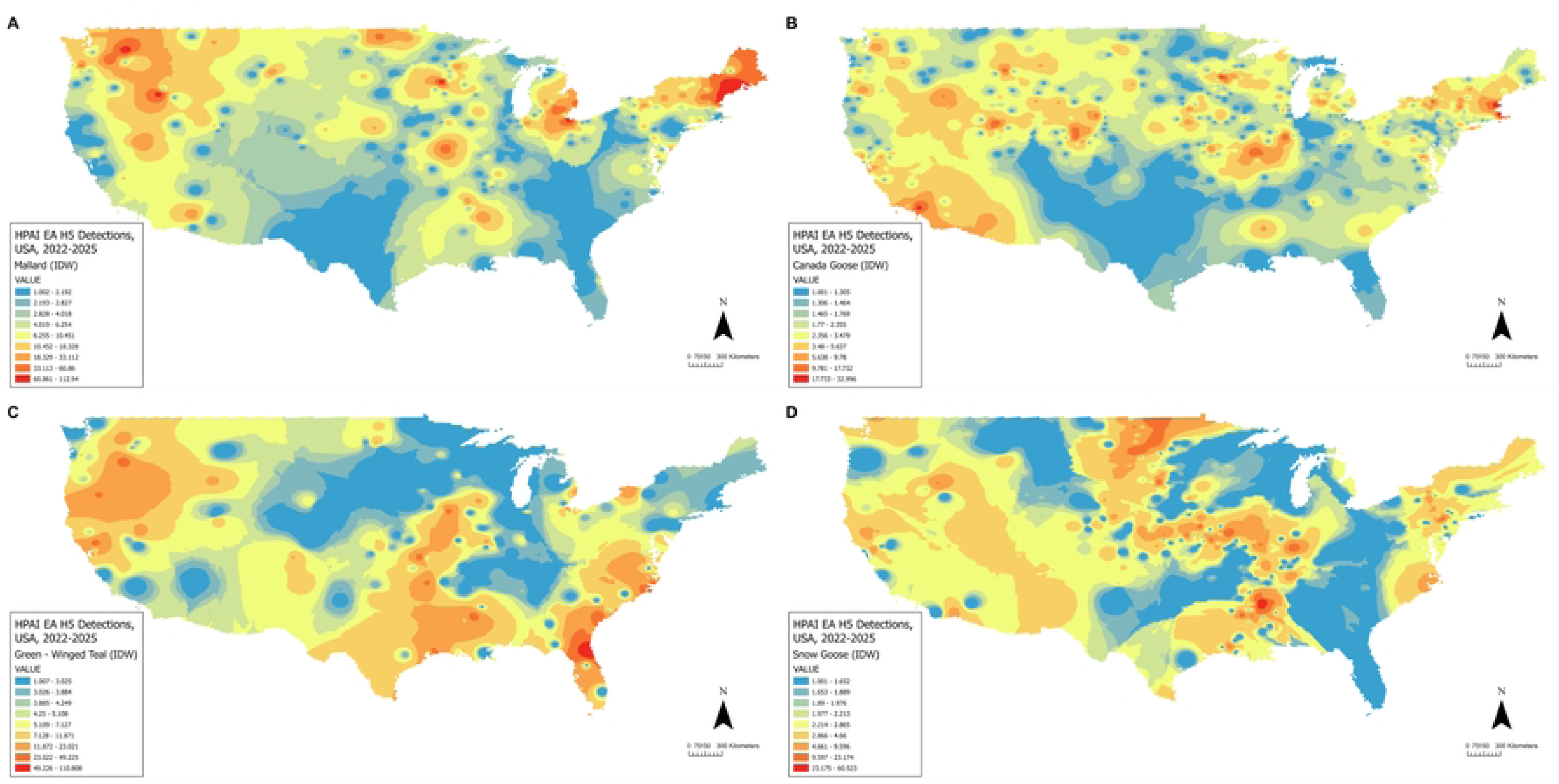
Spatial interpolation of highly pathogenic avian influenza (HPAI) H5 detections in wild waterfowl species using inverse distance weighting. (A) Mallard, (B) Canada goose, (C) Green-winged teal, and (D) Snow Goose. High detection areas are shown in red, medium detection areas in yellow tones, and low detection areas in blue.

### Space-time clustering of HPAI H5 among wild waterfowl

The space-time permutation scan statistic identified several significant space-time clusters for each wild bird species, where observed detections exceeded expected levels (O/E > 1). These clusters varied in magnitude, duration, and spatial extent, reflecting periods of elevated detection intensity (**Figure 3**, **Table 1**). Seven clusters were identified for Mallard. The primary cluster (O/E = 4.63, Feb-Mar 2025) occurred in the Northeast, while a high-intensity cluster (O/E = 14.42) centered on Minnesota during spring to early fall 2025. Other clusters were in Michigan (O/E = 8.73, Oct 2025) and the Pacific Northwest (O/E = 8.57, Nov 2022). Canada goose had seven clusters. The primary cluster (O/E = 3.77, May-Sept 2022) spanned the West Coast and Northwest. A high-intensity cluster (O/E = 30.16) occurred in the Midwest (Jan 2025). Additional clusters were in the Northeast and Pacific Northwest. Ten clusters were identified for Green-winged teal, with the primary cluster (O/E = 7.92) in Louisiana (Nov 2022). A high-intensity spike (O/E = 58.29, Jul 2024) was observed in Oregon. Other clusters spanned the Southeast, Midwest, and West Coast. Five Snow goose clusters were found, including a primary cluster (O/E = 6.60, Apr–May 2022) in the Northern Plains. A long-duration cluster (O/E = 2.06) spanned two years in the South, while the strongest cluster (O/E = 26.21) was centered in California during late winter 2024.

**Figure 3.**
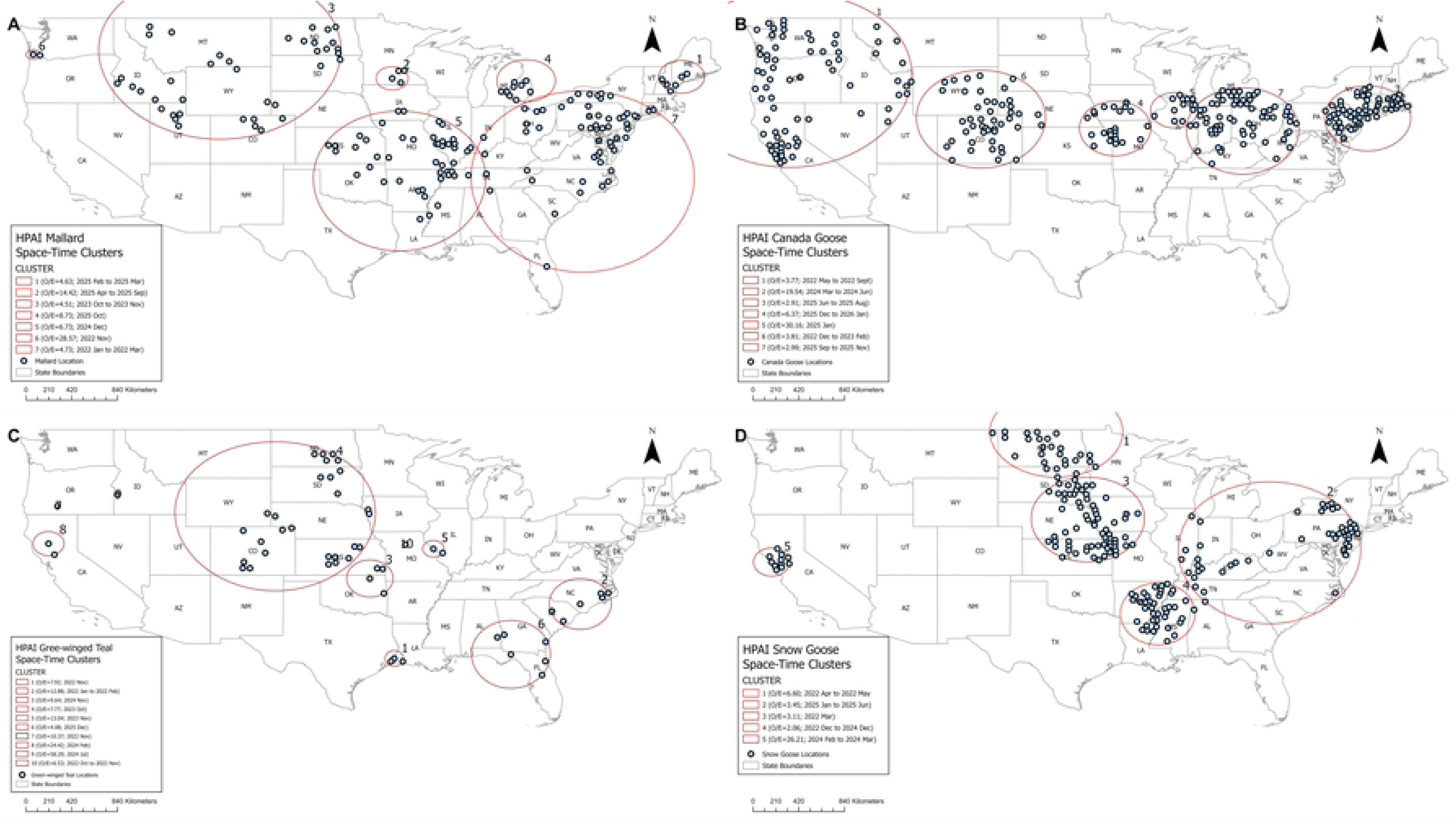
Results of space-time permutation models for highly pathogenic avian influenza (HPAI) H5 detections in wild waterfowl species across the U.S., 2022-2025. (A) Mallard, (B) Canada goose, (C) Green-winged teal, and (D) Snow Goose. The analysis used a cylindrical scanning window with 50% of the population and 50% of the study period, highlighting significant space-time clusters of higher than expected detections. Significant at p≤0.05, using 999 Monte Carlo simulations.

**Table 1.** Space-time clusters of highly pathogenic avian influenza (HPAI) H5 detections in major wild bird species, United States, 2022–2025. For each wild waterfowl species type, Cluster 1 represents the primary cluster (i.e., the most likely cluster with the highest test statistic), followed by secondary clusters in decreasing order of importance. Clusters were identified using the retrospective space-time permutation scan statistic with a scanning window maximum of up to 50% of the background population at risk and 50% of the study period. Statistical significance was evaluated using 999 Monte Carlo replications, with clusters considered significant at p ≤ 0.05. Reported radii represent the spatial extent of the detected clusters, and coordinates indicate the centroid of each identified cluster.

| Species | Cluster | Center coords<br>(Lat, Long) | Radius<br>(km) | Time frame | Cases | Expected<br>cases | Observed /<br>Expected_ | Test<br>statistic | P value |
| --- | --- | --- | --- | --- | --- | --- | --- | --- | --- |
| Mallard | 1 | 44.1658, -70.2065 | 150.38 | 2025-02 to 2025-03 | 222 | 48 | 4.63 | 171.5111 | 1.00E-17 |
|  | 2 | 44.0346, -94.0670 | 106.53 | 2025-04-01 to 2025-09-30 | 97 | 6.73 | 14.42 | 170.0266 | 1.00E-17 |
|  | 3 | 45.9373, -108.2744 | 769.72 | 2023-10 to 2023-11 | 186 | 41.24 | 4.51 | 139.2159 | 1.00E-17 |
|  | 4 | 43.8333, -83.0190 | 193.12 | 2025-10 | 103 | 11.79 | 8.73 | 133.4844 | 1.00E-17 |
|  | 5 | 35.5697, -93.4602 | 641.37 | 2024-12 to 2024-12 | 112 | 16.63 | 6.73 | 119.8432 | 1.00E-17 |
|  | 6 | 45.9950, -123.6557 | 44.19 | 2022-11 | 39 | 1.37 | 28.57 | 93.35412 | 1.00E-17 |
|  | 7 | 35.5178, -78.3657 | 830.49 | 2022-01 to 2022-03 | 72 | 15.54 | 4.63 | 54.51441 | 1.00E-17 |
| Canada goose | 1 | 43.9150, -121.2281 | 802.89 | 2022-05 to 2022-09 | 117 | 31.06 | 3.77 | 71.8141 | 1.00E-17 |
|  | 2 | 42.6732, -70.9524 | 0 | 2024-03 to 2024-06 | 27 | 1.38 | 19.54 | 54.86091 | 1.00E-17 |
|  | 3 | 40.7326, -73.5862 | 299.32 | 2025-06 to 2025-08 | 118 | 40.59 | 2.91 | 50.61219 | 1.00E-17 |
|  | 4 | 39.8932, -94.4047 | 250.85 | 2025-12 to 2026-01 | 37 | 5.81 | 6.37 | 37.63012 | 3.80E-15 |
|  | 5 | 41.4041, -89.5286 | 162.35 | 2025-01 | 15 | 0.5 | 30.16 | 36.66627 | 1.10E-14 |
|  | 6 | 40.6664, -105.4611 | 448.39 | 2022-12 to 2023-02 | 56 | 14.69 | 3.81 | 34.20265 | 1.90E-13 |
|  | 7 | 39.6915, -83.8899 | 396.17 | 2025-09 to 2025-11 | 55 | 18.4 | 2.99 | 24.08974 | 1.80E-08 |
| Green-winged teal | 1 | 30.1213, -93.8939 | 72.72 | 2022-11 | 49 | 6.19 | 7.92 | 59.25336 | 1.00E-17 |
|  | 2 | 34.6146, -78.5637 | 233.6 | 2022-01 to 2022-02 | 34 | 2.64 | 12.88 | 55.90216 | 1.00E-17 |
|  | 3 | 36.7152, -95.9044 | 170.36 | 2024-11 | 34 | 3.53 | 9.64 | 46.91715 | 1.00E-17 |
|  | 4 | 41.8506, -103.7080 | 683.02 | 2023-10 | 36 | 4.63 | 7.77 | 42.79614 | 1.00E-17 |
|  | 5 | 39.1693, -90.6676 | 75.91 | 2023-11 | 24 | 1.84 | 13.04 | 39.6492 | 1.00E-17 |
|  | 6 | 30.4580, -84.2779 | 311.48 | 2025-12 | 53 | 12.99 | 4.08 | 35.12683 | 6.30E-15 |
|  | 7 | 42.6864, -121.6501 | 0 | 2022-11 | 23 | 2.22 | 10.37 | 33.18113 | 5.90E-14 |
|  | 8 | 39.5981, -122.3919 | 109.77 | 2024-02 | 14 | 0.57 | 24.42 | 31.37496 | 4.80E-13 |
|  | 9 | 43.6251, -116.7093 | 0 | 2024-07 | 10 | 0.17 | 58.29 | 30.86146 | 8.60E-13 |
|  | 10 | 39.5151, -92.9626 | 0 | 2022-10 to 2022-11 | 20 | 3.06 | 6.53 | 20.68718 | 1.10E-07 |
| Snow goose | 1 | 48.6855, -99.2457 | 401.73 | 2022-04 to 2022-05 | 89 | 13.49 | 6.60 | 95.11838 | 1.00E-17 |
|  | 2 | 38.8345, -81.6748 | 650.02 | 2025-01 to 2025-06 | 102 | 29.55 | 3.45 | 56.46517 | 1.00E-17 |
|  | 3 | 41.5779, -96.6540 | 388.03 | 2022-03 | 89 | 28.59 | 3.11 | 42.41053 | 1.00E-17 |
|  | 4 | 33.7971, -90.8806 | 284.26 | 2022-12 to 2024-12 | 182 | 88.39 | 2.06 | 42.38332 | 1.00E-17 |
|  | 5 | 38.0735, -122.7236 | 129.75 | 2024-02 to 2024-03 | 17 | 0.65 | 26.21 | 39.29913 | 1.00E-17 |

When comparing wild waterfowl species, Mallard and Canada goose demonstrated broad, recurrent clusters spanning multiple flyways, whereas green-winged teal exhibited short-duration, localized, high-intensity bursts. Snow geese displayed fewer clusters but sustained low-to-moderate elevation across extended periods.

### Distribution of EA H5 HPAI detections in commercial poultry operations

Turkey operations had 532 outbreaks across 121 locations between February 8^th,^ 2022, and January 16^th,^ 2026 (**Figure 4**). A high number of detections were in the Upper Midwest (Minnesota and South Dakota), the Great Lakes (Michigan, Ohio), and the Intermountain West (Utah). Table-egg layer operations had 148 outbreaks across 55 locations between February 22^nd,^ 2022, and January 6^th^, 2026. A high number of detections were in the Midwest (Ohio, Indiana, Iowa), the Mountain West (Colorado, Arizona), the Mid-Atlantic (Pennsylvania, Delaware), and the West Coast (California). Broiler chicken operations exhibited 99 outbreaks across 54 locations between February 12^th,^ 2022, and January 13^th^, 2026. A high number of detections were in the West Coast (California), the Mid-Atlantic (Pennsylvania, Delaware), and the South (Arkansas, Tennessee). Commercial duck operations had 89 outbreaks across 20 locations between April 8^th^, 2022, and December 29^th^, 2025. A high number of detections were in the Midwest (Indiana, Ohio), the Mid-Atlantic (Pennsylvania), and the West Coast (California).

**Figure 4.**
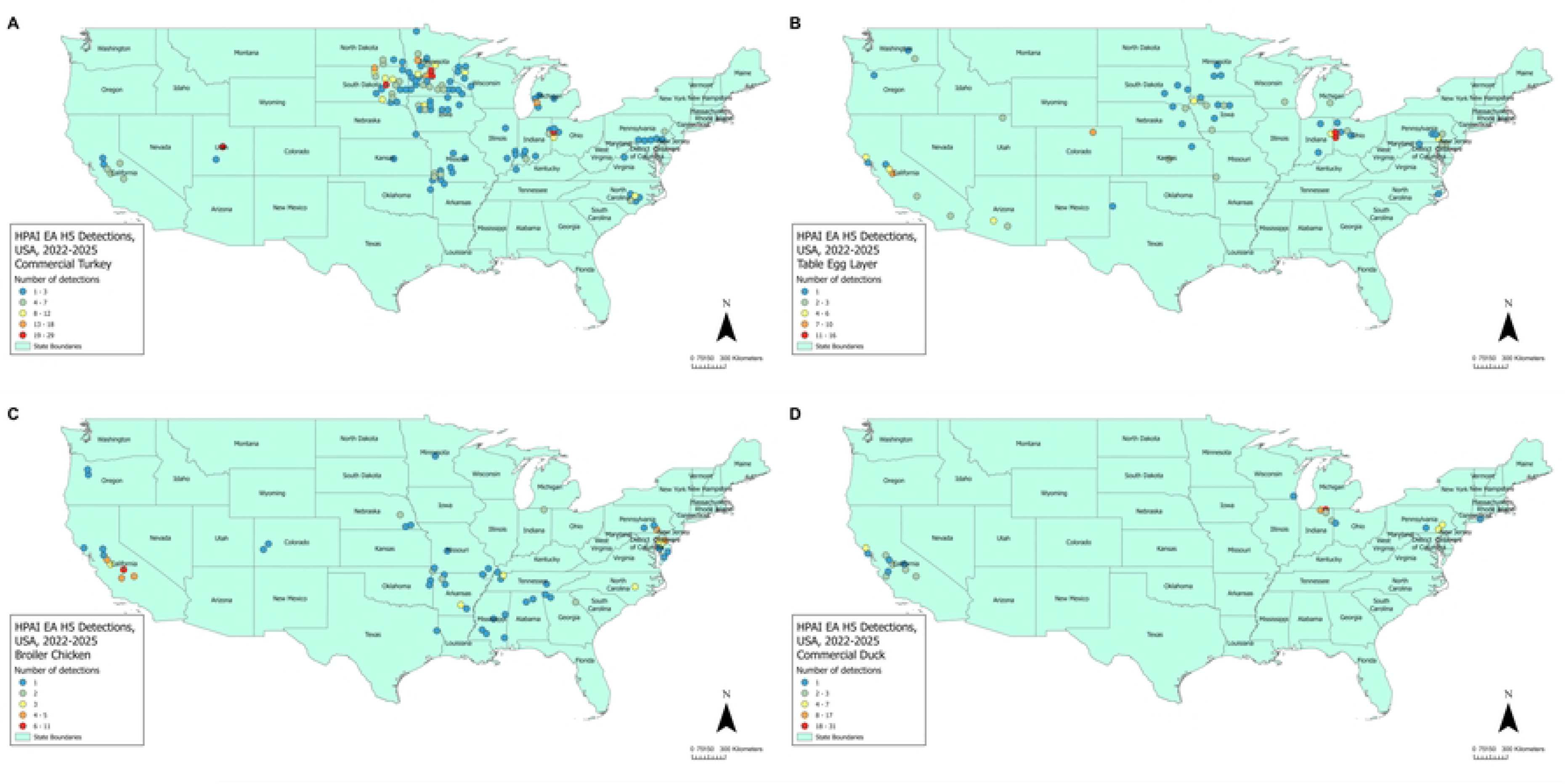
Distribution of highly pathogenic avian influenza (HPAI) H5 outbreaks in commercial poultry farms across the U.S., 2022–2025. (A)Turkey, (B)Table egg layer, (C) Broiler Chicken, and (D) Duck. Divergent colors are used, with red indicating high and blue indicating low outbreak number areas.

### Space-time clustering of HPAI H5 among commercial poultry operations

Across commercial poultry operations, multiple significant space-time clusters of HPAI (EA H5) were identified, varying by production type (**Figure 5**, **Table 2**). Commercial turkeys exhibited five clusters. Cluster 1 (O/E = 4.88; Jan–Mar 2025) occurred in the Upper Midwest. Cluster 2 (O/E = 7.31; Jul-Oct 2022) spanned the Upper Midwest and Northern Plains.

**Figure 5.**
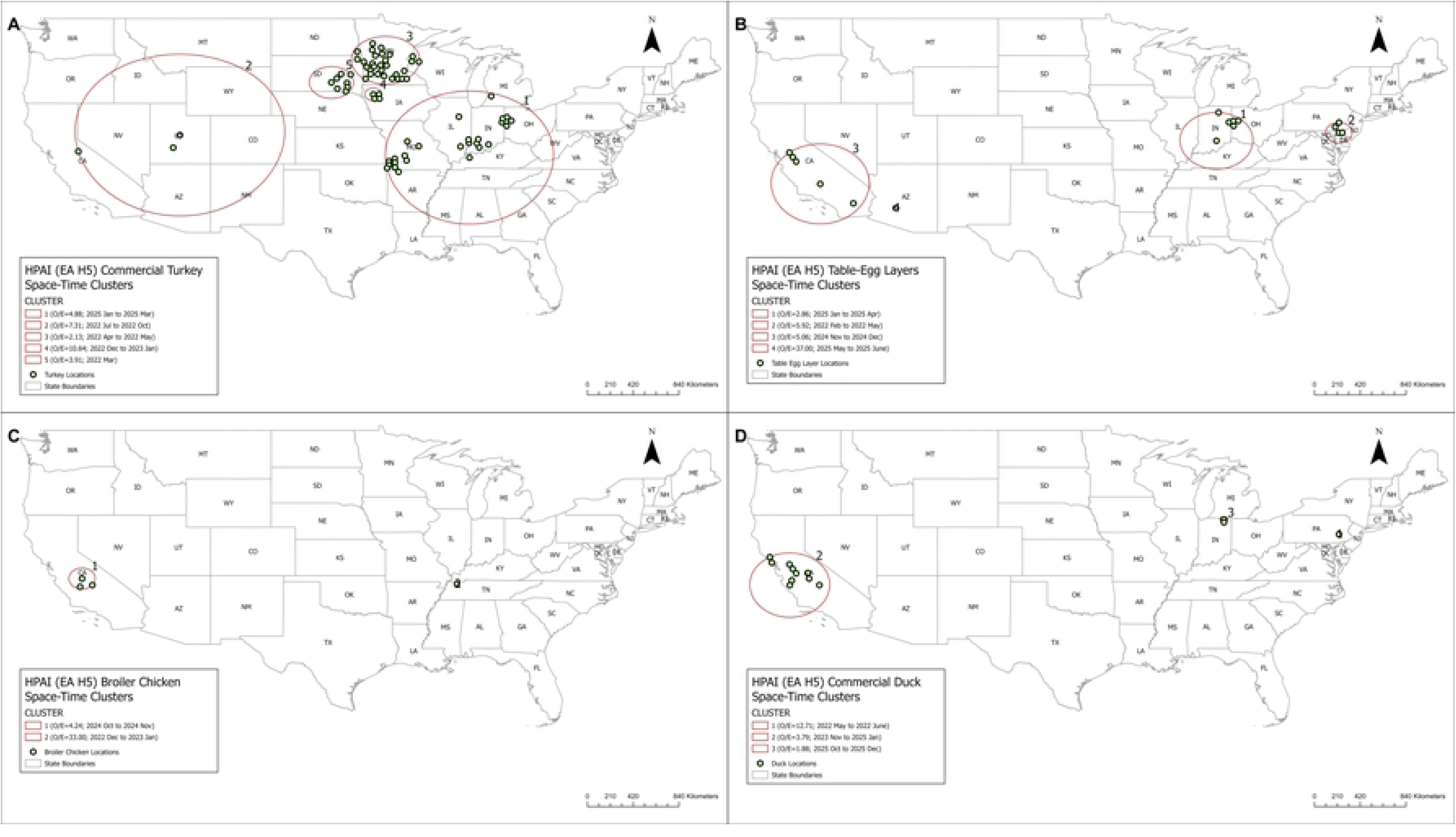
Space-time permutation model results for highly pathogenic avian influenza (HPAI) H5 outbreaks in poultry species across the U.S., 2022-2025. (A)Turkey, (B)Table egg layer, (C) Broiler Chicken, and (D) Duck. The analysis used a cylindrical scanning window with 50% of the population and 50% of the study period, highlighting significant space-time clusters of higher than expected outbreaks. Significant at p≤0.05, using 999 Monte Carlo simulations.

**Table 2.**
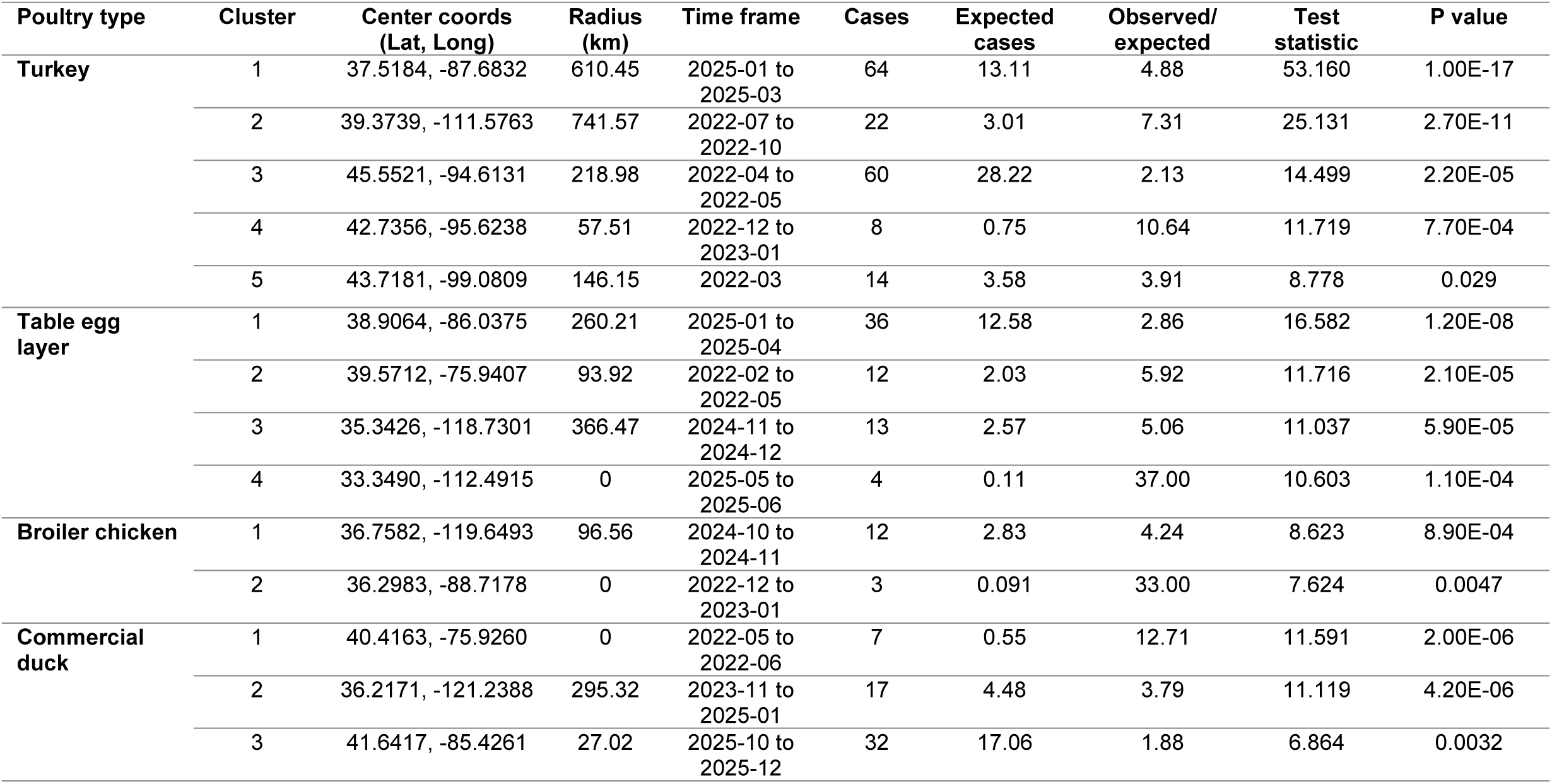
Space-time clusters of highly pathogenic avian influenza (HPAI) H5 outbreaks across major U.S. poultry sectors, 2022–2025. For each poultry type, Cluster 1 represents the primary cluster (i.e., the most likely cluster with the highest test statistic), followed by secondary clusters in decreasing order of importance. Clusters were identified using the retrospective space-time permutation scan statistic with a scanning window maximum of up to 50% of the background population at risk and 50% of the study period. Statistical significance was evaluated using 999 Monte Carlo replications, with clusters considered significant at p ≤ 0.05. Reported radii represent the spatial extent of the detected clusters, and coordinates indicate the centroid of each identified cluster.

| Poultry type | Cluster | Center coords<br>(Lat, Long) | Radius<br>(km) | Time frame | Cases | Expected<br>cases | Observed/<br>expected | Test<br>statistic | P value |
| --- | --- | --- | --- | --- | --- | --- | --- | --- | --- |
| Turkey | 1 | 37.5184, -87.6832 | 610.45 | 2025-01 to<br>2025-03 | 64 | 13.11 | 4.88 | 53.160 | 1.00E-17 |
|  | 2 | 39.3739, -111.5763 | 741.57 | 2022-07 to<br>2022-10 | 22 | 3.01 | 7.31 | 25.131 | 2.70E-11 |
|  | 3 | 45.5521, -94.6131 | 218.98 | 2022-04 to<br>2022-05 | 60 | 28.22 | 2.13 | 14.499 | 2.20E-05 |
|  | 4 | 42.7356, -95.6238 | 57.51 | 2022-12 to<br>2023-01 | 8 | 0.75 | 10.64 | 11.719 | 7.70E-04 |
|  | 5 | 43.7181, -99.0809 | 146.15 | 2022-03 | 14 | 3.58 | 3.91 | 8.778 | 0.029 |
| Table egg<br>layer | 1 | 38.9064, -86.0375 | 260.21 | 2025-01 to<br>2025-04 | 36 | 12.58 | 2.86 | 16.582 | 1.20E-08 |
|  | 2 | 39.5712, -75.9407 | 93.92 | 2022-02 to<br>2022-05 | 12 | 2.03 | 5.92 | 11.716 | 2.10E-05 |
|  | 3 | 35.3426, -118.7301 | 366.47 | 2024-11 to<br>2024-12 | 13 | 2.57 | 5.06 | 11.037 | 5.90E-05 |
|  | 4 | 33.3490, -112.4915 | 0 | 2025-05 to<br>2025-06 | 4 | 0.11 | 37.00 | 10.603 | 1.10E-04 |
| Broiler chicken | 1 | 36.7582, -119.6493 | 96.56 | 2024-10 to<br>2024-11 | 12 | 2.83 | 4.24 | 8.623 | 8.90E-04 |
|  | 2 | 36.2983, -88.7178 | 0 | 2022-12 to<br>2023-01 | 3 | 0.091 | 33.00 | 7.624 | 0.0047 |
| Commercial<br>duck | 1 | 40.4163, -75.9260 | 0 | 2022-05 to<br>2022-06 | 7 | 0.55 | 12.71 | 11.591 | 2.00E-06 |
|  | 2 | 36.2171, -121.2388 | 295.32 | 2023-11 to<br>2025-01 | 17 | 4.48 | 3.79 | 11.119 | 4.20E-06 |
|  | 3 | 41.6417, -85.4261 | 27.02 | 2025-10 to<br>2025-12 | 32 | 17.06 | 1.88 | 6.864 | 0.0032 |

Cluster 3 (O/E = 2.13; Apr-May 2022) reflected early spring activity in the central United States. Clusters 4 (O/E = 10.64; Dec 2022-Jan 2023) and 5 (O/E = 3.91; Mar 2022) represented overlapping winter hotspots in the Upper Midwest. Table-egg layers showed four clusters. Cluster 1 (O/E = 2.86; Jan-Apr 2025) occurred in Indiana and Ohio. Cluster 2 (O/E = 5.92; Feb-May 2022) was centered in Pennsylvania and Delaware. Cluster 3 (O/E = 5.06; Nov-Dec 2024) occurred in California, and Cluster 4 (O/E = 37.00; May-Jun 2025) represented an intense, short-duration cluster in Arizona. Commercial broilers had two clusters. Cluster 1 (O/E = 4.42; Feb-Apr 2024) occurred in California, while Cluster 2 (O/E = 6.89; Nov-Dec 2023) was centered in Tennessee, each reflecting localized periods of elevated activity. Commercial ducks exhibited three clusters. Cluster 1 (O/E = 12.71; May–Jun 2022) occurred in Pennsylvania. Cluster 2 (O/E = 3.79; Nov 2023–Jan 2025) was centered in California, and Cluster 3 (O/E = 1.88; Oct–Dec 2025) occurred in Indiana, marking a late-year cluster of lower relative intensity.

Spatiotemporal overlap between the primary turkey and layer outbreak clusters in the Midwest (Ohio and Indiana) in early 2025 was observed. In addition, spatiotemporal overlap between outbreaks in table egg layers (Cluster 3), broiler chickens (Cluster 1), and commercial ducks (Cluster 2) in the Pacific Coast region (California) during the fall of 2024 was identified. These overlaps occurred in high poultry production density regions in proximity to major migratory flyways.

### Spillover events of HPAI among wild birds and commercial poultry farms

Several spatiotemporal associations between wild waterfowl detections and poultry outbreaks were identified. The first spillover event occurred during winter-spring 2022 in the Northeast U.S. (Atlantic Flyway), where detections in Mallards (Cluster 7; January-March 2022) preceded outbreaks in table-egg layer (Cluster 2; February-May 2022) and commercial duck farms (Cluster 1; May-June 2022). The second event occurred during spring 2022 in the Upper Midwest (Mississippi Flyway), where snow goose detections (Cluster 3; March 2022; Cluster 1; April-May 2022) coincided with outbreaks in commercial turkey farms (Cluster 5; March 2022 and Cluster 3; April-May 2022). The third event occurred in the Pacific and Mountain West regions (Pacific Flyway), where detections in Canada goose (Cluster 1; May-September 2022) coincided with turkey outbreaks (Cluster 2; July-October 2022). The fourth event occurred during winter 2022-2023 in the Upper Midwest, where Canada goose detections (Cluster 6; December 2022-February 2023) in the Midwest and Mountain West overlapped with commercial turkey outbreaks (Cluster 4; December 2022-January 2023). The fifth event occurred during the fall-winter of 2025 in the Pacific region (Pacific Flyway), where snow goose (Cluster 5; February-March 2024) and green-winged teal (Cluster 8; February 2024) detections coincided with duck outbreaks (Cluster 2; November 2023–January 2025). Additionally, outbreaks in broiler chickens (Cluster 1; October-November 2024) and table-egg layers (Cluster 3; November-December 2024) were seen in this region. The sixth event occurred during the winter-spring of 2025 in the Midwest (Mississippi Flyway), where snow goose (Cluster 2; January-June 2025) and Canada goose (Cluster 5; January 2025) detections coincided with outbreaks in turkey (Cluster 1; January-March 2025) and table-egg layer (Cluster 1; January-April 2025) farms. The last spillover event occurred during the fall-winter of 2025 in the Upper Midwest, where Canada goose (Cluster 7; September-November 2025) and mallard (Cluster 4; October 2025) detections overlapped with outbreaks in commercial duck farms (Cluster 3; October-December 2025).

## Discussion

This study analyzed surveillance data on wild waterfowl HPAI H5 detections and commercial poultry outbreaks from February 2022 until early January 2026. Disease mapping and space-time permutation models analyzed wild waterfowl detections and commercial poultry outbreaks, and identified distinct regional and temporal patterns. The spatiotemporal analysis identified seven distinct spillover events, suggesting various HPAI H5 transmission pathways at the wild waterfowl-commercial poultry farm interface. These events can be categorized into four phases. The first phase covered winter-spring 2022, corresponding to the initial incursion of the virus via the Atlantic Flyway [11], and the amplification of the virus in Mallards before spillover to commercial ducks and table-egg layers (Event 1). The second phase included the spring-fall 2022 period, when the virus expanded westward (Events 2 & 3) via the Mississippi Flyway, driven by the high viral environmental load from migration of Snow goose and Canada goose, where commercial turkey farms were affected. Phase 3 included the period between winter 2022 and 2024, where persistence of the virus in Canada goose and outbreaks in turkey farms occurred in the Midwest (Event 4). This phase also included the fourth event in the Pacific region, with a prolonged outbreak in commercial ducks showing a persistence of the virus (fall 2023 to winter 2025) and short localized outbreaks in broiler chickens and table egg layers in fall 2024. In this region, clusters were identified in Snow goose and Green-winged teal during winter 2024, suggesting an expansion of the host range. Finally, Phase 4 included the winter of 2025, when the virus resurged in the Midwest (Events 6 & 7), suggesting a cyclical reinfection of the Mississippi Flyway, and high viral load and detections in Canada goose, Snow goose, and Mallard, and large outbreaks in turkey, table egg, and commercial duck farms, mirroring Phase 1 and 2 in its extent and severity.

This study identified vector-specific transmission dynamics. Canada geese were involved in the sustained persistence and spread of the virus to commercial turkey farms across the Midwest and Great Plains (Events 3, 4, 6, and 7). Canada goose populations are abundant in agricultural fields and in proximity to poultry farms [18], graze on farm peripheries and overwinter near human habitat, and might act as a local environmental reservoir and a bridge species, linking the gap between seasonal migration waves, allowing the virus to overwinter. In addition, in late 2025 (Event 7), the overlap of Canada goose detections (Cluster 7) with outbreaks in commercial duck farms suggests a transmission shift.

Our analysis of Event 2 (Spring 2022) and Event 6 (Winter-Spring 2025) identified a link between detections in Snow geese and outbreaks in commercial turkey operations. Snow geese are an effective HPAI H5 transmitter [19] migrating in large flocks, which facilitates rapid viral amplification. The detection of HPAI H5 in Snow geese (Clusters 1 and 3) immediately preceding outbreaks in Midwestern turkey farms suggests that high environmental viral loads facilitated viral transmission to turkey farms. The recurrence of this specific Goose-to-Turkey spillover pathway in 2025 in the Midwest underlines the vulnerability of turkey operations, especially during the winter and fall migration seasons, and calls for enhanced biosecurity programs [20]. Mallard detections were associated with outbreaks in commercial duck and layer farms (Events 1 and 7), suggesting that they act as maintenance hosts. Previous studies described that Mallards can be infected and shed high viral loads, but they remain asymptomatic or exhibit only mild clinical signs [21]. This phenomenon poses a high risk for the commercial duck sector, where proximity to wild reservoirs can facilitate viral amplification and transmission if farm biosecurity breaches occur. The recurrence of this pattern in the Upper Midwest in late 2025 (Event 7) underscores the long-term risk posed by resident or semi-migratory Mallard populations overwintering near poultry production sites.

Within commercial poultry, turkeys and table-egg layers exhibited the most consistent and overlapping outbreak clusters, particularly in the Midwest. The recurrent co-localization of turkey and layer outbreaks in this region implies that these regions within major migratory flyways are at heightened risk and could function as amplification zones following wild-bird introductions. The Mississippi Flyway, home to high densities of commercial poultry in the Midwest, saw significant spillover events during the spring and fall migrations of 2022 and 2025. On the other hand, broiler and commercial duck clustering appeared more episodic or geographically focal.

Our analysis suggests that the current HPAI H5 strain, especially the 2.3.4.4b clade, can persist in the environment and can be sustained by the local wild waterfowl populations, posing a continuous risk to poultry operations [9]. The current outbreak has evolved from a seasonal, migratory introduction (2022) into a persistent, multi-vector endemic system (2025), with Snow Geese and Canada Geese emerging as the primary bridge vectors for commercial turkey flocks. Rather than a single introduction wave, the findings of our study support a pattern of repeated seasonal spillover events [22], followed by regional amplification within high-density poultry production systems, especially turkey and table egg layer farms, suggesting a possible spread via fomites.

Risk-based resource allocation, considering the location, poultry production types, and wild bird types, may improve outbreak detection and response efficiency compared with uniform national approaches. The recurrent overlap between turkey and layer clusters indicates that multi-commodity coordination in regions with active wild-bird pressure could improve outbreak control. Additionally, farm biosecurity programs should be updated and synchronized across poultry sectors, considering the local disease pressures, and they are cost-effective in preventing outbreaks [23]. Furthermore, implementing spatial-temporal cluster detection methods into surveillance systems [24] and integrating wild bird and poultry HPAI H5 surveillance could improve outbreak preparedness and control. Finally, linking migratory waterfowl detection intensity with poultry density maps could inform animal health authorities to anticipate risk and proactively implement prevention and control programs.

## Data Availability Statement

The data presented in this article is publicly available.

## Acknowledgment

I acknowledge the U.S. Department of Agriculture (USDA) Animal and Plant Health Inspection Service (APHIS) staff for collecting the samples and providing access to the HPAI H5 dataset.

## References

1. Spackman E. A brief introduction to avian influenza virus. Methods Mol Biol. 2020;2123:83–92. doi:10.1007/978-1-0716-0346-8_7

2. Bi Y, Yang J, Wang L, Ran L, Gao GF. Ecology and evolution of avian influenza viruses. Curr Biol. 2024;34(15):R716–R721. doi:10.1016/j.cub.2024.05.053

3. Damodaran L, Jaeger AS, Moncla LH. Ecology and spread of the North American H5N1 epizootic. Nature. 2026;649(8096):432–441. doi:10.1038/s41586-025-09737-x

4. Blagodatski A, Trutneva K, Glazova O, et al. Avian influenza in wild birds and poultry: Dissemination pathways, monitoring methods, and virus ecology. Pathogens. 2021;10(5):1–23. doi:10.3390/pathogens10050630

5. Couty M, Guinat C, Fornasiero D, et al. The role of wild birds in the global highly pathogenic avian influenza H5 panzootic, 2020–2023. NPJ Biodivers. 2026;5(1). doi:10.1038/s44185-025-00114-5

6. Caserta LC, Frye EA, Butt SL, et al. Spillover of highly pathogenic avian influenza H5N1 virus to dairy cattle. Nature. 2024;634(8034):669–676. doi:10.1038/s41586-024-07849-4

7. Tawidian P, Torchetti MK, Killian ML, et al. Genotypic Clustering of H5N1 Avian Influenza Viruses in North America Evaluated by Ordination Analysis. Viruses. 2024;16(12):1–17. doi:10.3390/v16121818

8. U.S. Department of Agriculture (USDA) Animal and Plant Health Inspection Service (APHIS). Confirmations of Highly Pathogenic Avian Influenza in Commercial and Backyard Flocks. https://www.aphis.usda.gov/livestock-poultry-disease/avian/avian-influenza/hpai-detections/commercial-backyard-flocks. Accessed January 10, 2026.

9. Fang K, Li J, Zhao H, et al. Assessing HPAI-H5 transmission risk across wild bird migratory flyways in the United States TI IN ES ES. Nat Commun. 2026. doi:10.1038/s41467-026-69344-w

10. U.S. Department of Agriculture (USDA) Animal and Plant Health Inspection Service (APHIS). Implementation Plan for Avian Influenza Surveillance in Waterfowl in the United States, Summer 2025 – Spring 2026.

11. Prosser DJ, Kent CM, Sullivan JD, et al. Using an adaptive modeling framework to identify avian influenza spillover risk at the wild-domestic interface. Sci Rep. 2024;14(1):1–13. doi:10.1038/s41598-024-64912-w

12. Prosser DJ, Chen J, Ahlstrom CA, et al. Maintenance and dissemination of avian-origin influenza A virus within the northern Atlantic Flyway of North America. PLoS Pathog. 2022;18(6):1–25. doi:10.1371/journal.ppat.1010605

13. U.S. Department of Agriculture (USDA) Animal and Plant Health Inspection Service (APHIS). Early Detection and Monitoring for Avian Influenzas of Significance in Wild Birds: A U.S. Interagency Strategic Plan, 2015.; 2015.

14. U.S. Department of Agriculture (USDA) Animal and Plant Health Inspection Service (APHIS). National List of Reportable Animal Diseases and Reporting Requirements for Highly Pathogenic Avian Influenza. https://www.aphis.usda.gov/livestock-poultry-disease/surveillance/reportable-diseases. Accessed January 8, 2026.

15. U.S. Department of Agriculture (USDA) Animal and Plant Health Inspection Service (APHIS). Detections of Highly Pathogenic Avian Influenza in Wild Birds. https://www.aphis.usda.gov/livestock-poultry-disease/avian/avian-influenza/hpai-detections/wild-birds. Accessed January 4, 2026.

16. R Core Team. R: A language and environment for statistical computing: R Foundation for Statistical Computing. https://www.r-project.org/.

17. Kulldorff M. SaTScanTM: Software for the spatial and space-time scan statistics. www.satscan.org.

18. Jimenez C, Kolokotronis SO, Rosenbaum JE, Hoepner LA. Evaluating the Role of Canada Goose Populations in Transmission Dynamics During Peak HPAI Incidence in Iowa, February 2022–December 2023. Appl Sci. 2025;15(12):1-17. doi:10.3390/app15126900

19. Sullivan JD, Casazza ML, Poulson RL, et al. Potential impacts of 2.3.4.4b highly pathogenic H5N1 avian influenza virus infection on Snow Goose (Anser caerulescens) movement ecology. PLoS One. 2025;20(7 July):1-15. doi:10.1371/journal.pone.0328149

20. Kang M, Wang LF, Sun BW, et al. Zoonotic infections by avian influenza virus: changing global epidemiology, investigation, and control. Lancet Infect Dis. 2024;24(8):e522–e531. doi:10.1016/S1473-3099(24)00234-2

21. Teitelbaum CS, Masto NM, Sullivan JD, et al. North American wintering mallards infected with highly pathogenic avian influenza show few signs of altered local or migratory movements. Sci Rep. 2023;13(1):1–11. doi:10.1038/s41598-023-40921-z

22. Russell SL, Andrew CL, Yang KC, et al. Descriptive epidemiology and phylogenetic analysis of highly pathogenic avian influenza H5N1 clade 2.3.4.4b in British Columbia (B.C.) and the Yukon, Canada, September 2022 to June 2023. Emerg Microbes Infect. 2024;13(1):1–13. doi:10.1080/22221751.2024.2392667

23. Koppes P, Guerrant T, Marks D, et al. An economic evaluation of preventing vs. suppressing HPAI outbreaks: A case study from Iowa. Prev Vet Med. 2025;244(February):106651. doi:10.1016/j.prevetmed.2025.106651

24. Agrawal I, Sharma B, Varga C. Space-Time Clustering and Climatic Risk Factors for Lumpy Skin Disease of Cattle in Uttar Pradesh, India, 2022. Transbound Emerg Dis. 2024;2024. doi:10.1155/2024/1343156

